# Facial color properties affect perceived age

**DOI:** 10.64898/2026.09.29.743707

**Authors:** Jean Hsieh, Colin W.G. Clifford, Erin Goddard

## Abstract

Age is a fundamental social attribute that affects rights, responsibilities and even social status. Color information of the face can affect perceived age, but the nature of its contribution, independent of luminance information, remains underexplored. To better characterize the role of color in age perception, we conducted two experiments. In our first experiment, we tested whether natural age-related changes in facial color alone can drive changes in perceived age. In our second experiment, we manipulated red-green contrast across faces aged 18-91 and tested the effect of this manipulation of perceived youthfulness. Our findings demonstrate that natural age-related color changes do provide cues for age perception, dissociable from luminance information. We also found that the optimal red-green contrast level to make faces look the youngest was significantly higher than the originals, meaning that faces tended to appear younger when their redness was accentuated, even though average face redness increases with age. Our analysis revealed that younger faces required greater increases in redness level compared to their older counterparts to achieve the youngest appearance. These results suggest that the spatial distribution of redness contributes to apparent youthfulness, especially in younger faces.

**Precis:** We find that facial color, independent of luminance information, carries meaningful cues to perceived age. Boosting red-green contrast of face images makes them appear younger, despite the fact that this increases their overall redness, which tends to increase with age.

---

Age is a crucial social attribute that influences rights, responsibilities and social status (Mayes et al., 2010a; Rhodes, 2009). We spontaneously judge the age of new acquaintances and strangers without realising we are doing so (Palermo & Rhodes, 2007). The proficiency in age estimation is linked to more socially appropriate interpersonal behaviour. For example, estimating whether a customer is old enough to buy age-restricted products like alcohol or tobacco is one of the legal uses for age estimation. Willner and Rowe (2001) examined the accuracy of 100 alcohol servers in estimating the ages of individuals aged 13, 16, 20 or 22, in a context with a legal drinking age of 18 years old. Participants significantly overestimated the ages of 13 and 16 year-olds: 38% of 16-year-old boys and 56% of 16-year-old girls were judged old enough to buy alcohol, highlighting the possibility of unintentional alcohol sales to minors resulting from age misperceptions (Willner & Rowe, 2001). Despite the high social and legal importance of age perception, several factors remain unknown in how different facial cues contribute to the perception of age.

The process of age estimation heavily relies on faces, as faces stand out as a reliable and robust source of age-related information throughout an individual’s lifetime (Rhodes, 2009). Several candidate facial cues have been suggested as relevant, which can be broadly categorized into feature cues, configural cues and surface cues (Bruce & Langton, 1994; Diamond & Carey, 1986; George & Hole, 1998, 2000; Rhodes, 2009). Feature cues involve internal facial components, such as eyes and lips. The eyes and lips have been reported to be particularly relevant to estimating age compared to other facial features. The surrounding area, shape, and size of the eyes and sclera color have been shown to help people make a judgement of age (Russell et al., 2014). Lips are also potentially a reliable cue to age since their volume (Gunn et al., 2009) and redness (Gomi & Imamura, 2019) decrease significantly with age. In addition, spatial relationships between features, like the distance between the mouth and nose, have been proposed as configural cues for age estimation (George & Hole, 2000; termed ’global features’ in Rhodes, 2009).

Compared with feature and configural cues, surface-based cues to perceived age remain less explored. Although the term ‘surface-based’ most naturally encompasses variation due to skin pigmentation, tone and texture, the three-dimensional (3D) structure of faces can create patterns of shading and shadows in images of faces that could also provide cues to age, and these are usually conflated with skin properties in natural images of faces under typical lighting conditions. For this reason, surface-based cues associated with perceived age have been functionally divided into two categories (Bruce & Langton, 1994; Kemp et al., 1996; Russell et al., 2006). The first category, known as “pigmentation,” includes variations in skin tone and texture (Bruce & Langton, 1994). Facial skin dyspigmentation (Dobos et al., 2015) and the number and size of pigmented spots (Kikuchi et al., 2022) increase with age, and faces with uneven skin tone and hyperpigmentation are more likely to be judged as older compared to those with even skin tone (Perrett, Bunse, et al., 2012; Gunn et al., 2009; Mayes et al., 2010b). Sun exposure is also associated with a higher perceived age (Leyden, 1990; Mayes et al., 2010b; Ortonne, 1990) and more uneven skin tone (Matts et al., 2007). The second category is the pattern of shading and shadow created by the 3D shape of the face (Bruce & Langton, 1994; Hill et al., 1995; Kemp et al., 1996). For example, the shading and shadowing produced by the nose’s 3D shape changes significantly with age (Sforza et al., 2011), but it is not yet clear if people use this cue to estimate age. Ageing and skin sun-damage are both associated with wrinkling (Gunn et al., 2009). Empirically, digitally reducing skin wrinkles, skin sagging, or skin unevenness has been shown to give a more youthful appearance (Fink et al., 2006; Matts et al., 2007).

We are interested in characterising the role of color information in affecting the perceived age of faces. We have previously demonstrated that presenting face images in color rather than matched-luminance greyscale reduces inter-observer variability in rated ages, suggesting that color information is used in age estimation (Hsieh et al., 2023). We have also shown, for images of young adults, that increasing the contrast of isoluminant red-green information of faces alters perceived age, with higher-contrast, redder images tending to be rated younger. However, this effect is unlikely to be as simple as average redness increasing apparent youthfulness, since studies testing the correlation between natural inter-stimulus variation in average redness and perceived age have been inconsistent: Lu et al. (2021) found that perceived age was significantly positively correlated with in average redness for Caucasian face stimuli aged 20-40 years, but Voegeli et al. (2023) found a significant effect in the opposite direction for Caucasian face stimuli aged 20-69 years (neither study found an significant association between average redness and perceived age for other ethnic groups). These mixed results suggest that average redness alone is unlikely to account for the effect of increasing the contrast of red-green contrast (hereafter referred to as RG contrast) on perceived age.

Previous work has demonstrated that manipulating surface information for stimuli with identical spatial structure can alter perceived age by 5-20 years (Burt & Perrett, 1995; Fink et al., 2006). Fink et al. (2006) applied skin color distributions from different ages to an identical 3D shape-standardized model of a female face, and asked participants to rate the age, health, and attractiveness of the 3D rendered artificial faces. Faces with younger surface-based information were perceived as younger and received significantly higher ratings for attractiveness and health. Fink et al. (2006) note that skin color distributions tend to become less even with age, an observation confirmed with quantitative analyses showing that facial skin heterogeneity, measured within a patch of skin on the cheek, increases with age (Fink, Matts, et al., 2012; Matts et al., 2007) and perceived age (Nkengne et al., 2008). These results suggest a role for surface cues in perceived age, but they do not isolate the role of chromatic information from achromatic information.

Color information could also contribute to ‘facial contrast’, which is defined as the contrast between facial features such as the eyes and lips and the surrounding skin, and is another potential cue to age (Porcheron et al., 2013; Russell, 2003; Russell et al., 2017). Note that ‘facial contrast’, or the contrast between facial features and surrounding skin, is different from ‘skin contrast’, or the contrast within the skin surface, reflecting its inhomogeneity, as defined by others (Matts et al., 2007). Porcheron et al. (2013) proposed that decreased facial contrast with age serves as a cue for identifying age. They measured the facial contrast of Caucasian female faces within the CIE L*a*b* color space and asked participants to estimate age from the original and contrast-manipulated images. Female face images were judged to be significantly younger when shown with greater a* (red-green) contrast around the mouth, greater luminance contrast around the eyes, or greater luminance contrast around the eyebrows. However, these estimates of facial contrast do not distinguish between skin pigmentation and luminance cues related to topographic factors, and measuring facial contrast requires individual labelling of facial regions of interest, which makes standardization difficult. Overall, these findings suggested that color contrast, both between features and the skin and between features, is a potential cue to age.

A key challenge for facial age estimation research is the complexity of faces as stimuli, meaning that the number of possible variables driving age perception is large. A common approach is to manipulate specific facial features and observe changes in age ratings to identify relevant features (Fink et al., 2006; Porcheron et al., 2013). While this approach is valid, it relies on the researcher first identifying candidate facial features, leaving open the possibility that more complex or subtle features are missed. To circumvent this limitation, and to isolate the role of color information from luminance-driven cues, we manipulate the color contrast of entire real face images using the three dimensions of CIE L*a*b* space, as opposed to manipulating specific facial features.

Across two experiments, we tested how perceived age was affected when we globally manipulated the isoluminant color content of images of faces across adulthood (aged 18-91 years). In the first experiment, we show that faces with identical luminance content are perceived as older when combined with an isoluminant mask based on older faces, than when the isoluminant content is based on younger faces. In the second experiment, we asked participants to select the RG contrast level that made a series of faces appear youngest. Despite the fact that face overall redness tends to increase with age, participants consistently selected images with increased RG contrast (which have increased redness) as the most youthful in appearance. Our results demonstrate that isoluminant color information contributes to perceived age, and suggest that the spatial distribution of redness of faces influences the appearance of youthfulness.

## Stimuli

Stimuli for both Experiment 1 and 2 were based on face images sampled from the Park Aging Mind Laboratory Face Database (PFD) (Minear & Park, 2004), which includes self-reported race, gender and age at the time of the photograph. All selected images showed models photographed with neutral facial expressions who were self-identified Caucasians, and there were 80 male and 80 female models, ranging in age from 18 to 91. An additional 20 images from younger faces (aged 19 to 29) and 20 images from older faces (aged 70 to 93) were selected to generate the isoluminant masks used in Experiment 1.

### Image redness analysis

To characterize the age-related variation in face redness in our stimulus set, we measured the distributions of a* (red-green) values in each of the 160 base images used here, summarized in **Figure 1**. For each image, we considered only pixels within an ellipse-shaped face area (excluding the hair and background, described in detail in the methods for Experiment 1 below and illustrated in **Figure 2C**). We converted the image to CIE L*a*b* space, and considered only the a* values of each pixel. Across all 160 images, the a* values were bimodally distributed, with a dominant peak at around a*=20, and a smaller peak centred on approximately a*=0, as shown in the combined distribution in **Figure 1B**. To extract measures of dominant peak of a*, we modelled each images’ observed distribution of a* values with a mixture of 5 Gaussian components, estimated using the MATLAB function *fitgmdist.* To exclude the smaller peak centred on approximately a*=0, we removed Gaussian components with mean a*<5, using the remaining components as a model of the dominant peak of the a* values for the specified image.

**Figure 1:**
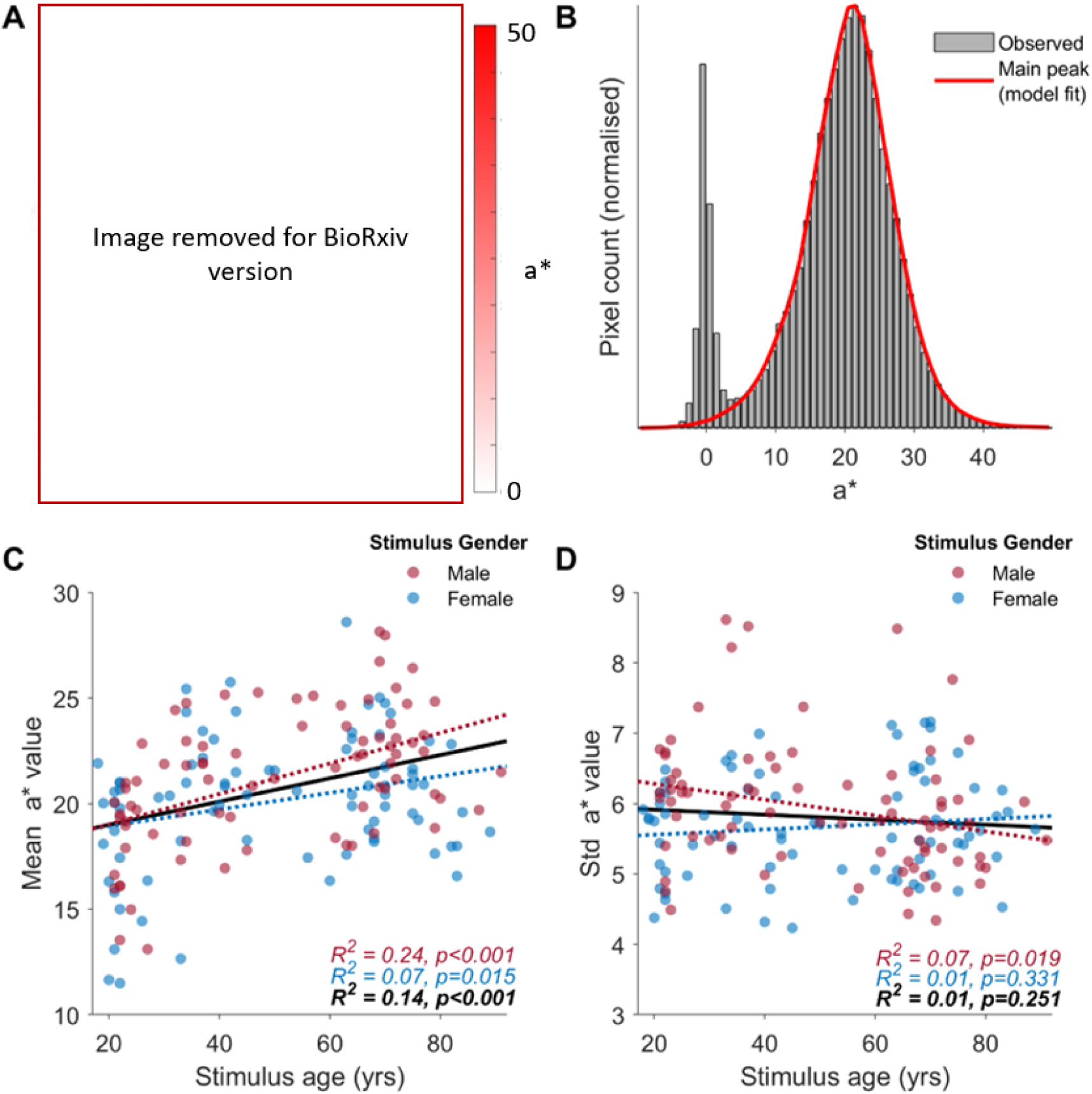
Red-green (a*) stimulus properties. **A**: Red-green (a*) value for 4 example stimuli. Here, and in subsequent figures, the eye regions are masked to conceal face identity, but these were unobscured in all stimulus images. **B**: Distribution of a* values across the 160 images from the Park Dataset used experiments, normalized so that each image contributes equally to the distribution. The red line shows the combined model distributions that were used to estimate the mean and standard deviation of the main peak of each images’ a* values (see text for details of modelling procedure). **C**-**D**: the mean (**C**) and standard deviations (**D**) of each image’s a* values, shown as a function of stimulus age and gender. Fitted lines and correlation coefficients are shown separately for the 80 male (red) and 80 female (blue) faces, and for the combined set of 160 faces (black).

**Figure 2:**
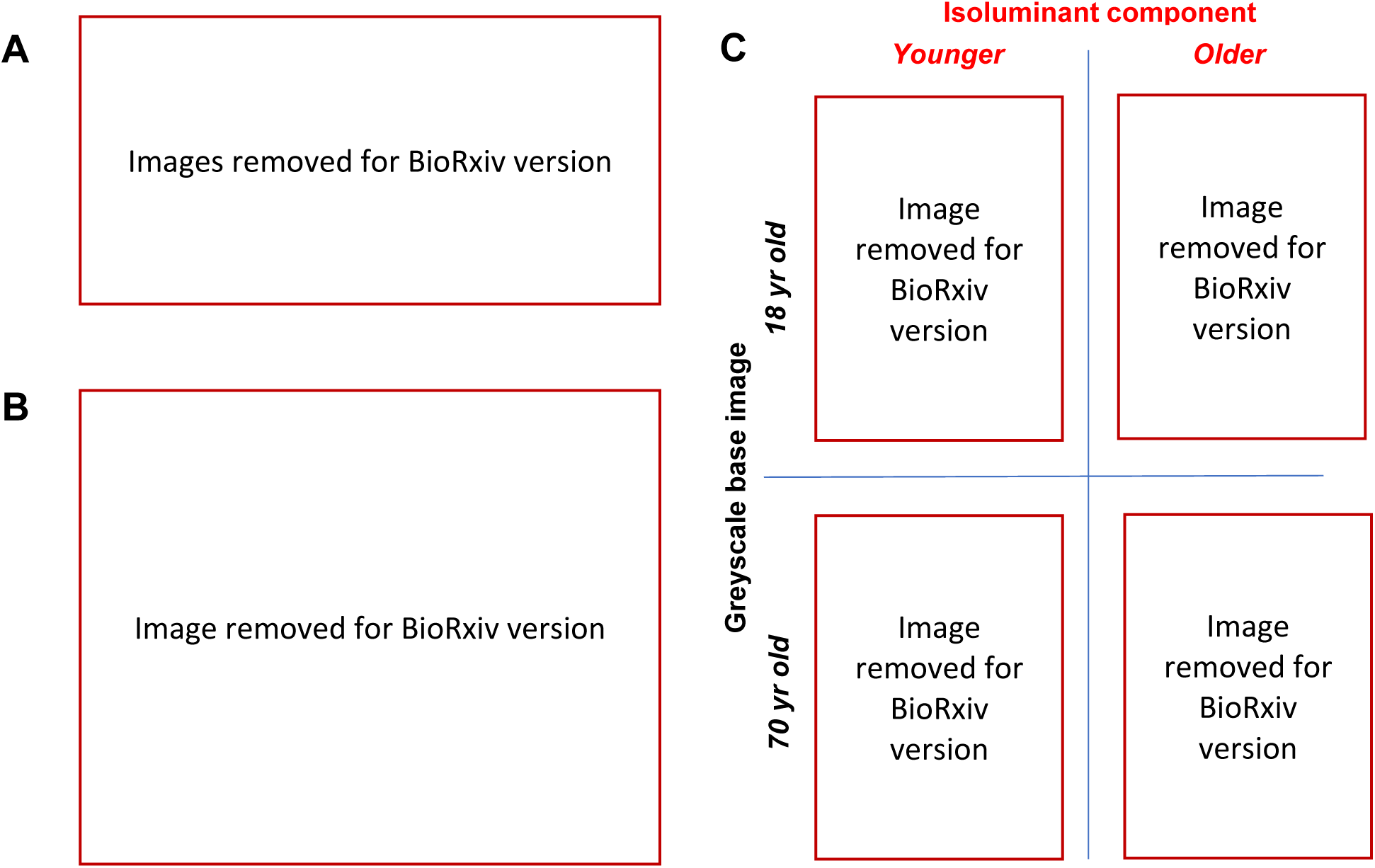
Stimulus Generation (Experiment 1). **A**: Original images were decomposed into isoluminant and greyscale components. **B**: 63 facial landmarks, shown here as red circles on an example face, were used to morph the isoluminant component of a younger or older image to the greyscale component of the base image. **C**: Example stimuli (composite faces) showing two greyscale base images (18 yr old and 70 yr old), each with a younger and older isoluminant component.

In **Figure 1C-D** we report the mean and standard deviation of these model distributions, as a function of stimulus age. For both male and female faces, there was a significant positive correlation between age and mean a* value of the dominant peak (male faces: R^2^=0.24, p<0.001; female faces R^2^=0.07, p=0.015; combined: R^2^=0.14, p<0.001). That is, the average a* value of the face images tends to increase in redness with age, but age was a fairly modest predictor of average a* value, accounting for 24% of the variance in average a* values for female faces, and just 7% of this variance for male faces. Standard deviations of a* values for each image were significantly negatively correlated with age for male faces (R^2^=0.07, p=0.019), but were not significantly correlated with age for female faces (R^2^=0.01, p=0.331), or for the combined set of images (R^2^=0.01, p=0.251).

We performed equivalent analyses on the distributions of b* (blue-yellow) values of each image (data not plotted), and found similar trends for ‘yellowness’ as for ‘redness’, consistent with both measures reflecting a common dimension with average skin hue intermediate to red and yellow. Specifically, there was a significant positive correlation between age and mean b* value (male faces: R^2^=0.27, p<0.001; female faces R^2^=0.07, p=0.022; combined: R^2^=0.15, p<0.001) but no significant correlations between age and the standard deviation of b* values in each image (male faces: R^2^=0.01, p=0.336; female faces R^2^=0.00, p=0.805; combined: R^2^=0.01, p=0.373).

### Experiment 1

#### Overview

In Experiment 1, we aimed to understand whether age-related changes in face color affect age perception. Applying skin color information from different ages to identical 2D or 3D shape-standardized faces has been shown to contribute to how we perceive the age of the face (Burt & Perrett, 1995; Fink et al., 2006). However, these previous studies did not exclude luminance information, making it difficult to attribute the effects solely to color. To obtain faces with purely chromatic information, we decomposed original color images into isoluminant color and greyscale components (see **Figure 2A**). The resulting isoluminant images were purely chromatic and did not have any luminance information. In Experiment 1, we extracted isoluminant color information from both young (age range from 19 – 29) and old (age range from 70 – 93) age groups and morphed them onto a new set of greyscale images across ages. Participants were asked to rate the ages of these composite faces. We hypothesized that a significant effect would be observed between conditions such that purely chromatic information would bias perceived age.

#### Methods

##### Participants

55 participants completed Experiment 1. After 3 exclusions (described under ‘Data cleaning’ below), there were 20 male, 30 female, and 2 other participants, aged 18-49 years (mean = 22.6, SD = 7.43). All participants had normal or corrected to normal vision and normal color vision (self-reported). All participants provided informed consent and ethical approval for this study was granted by the UNSW Human Research Ethics Committee. All methods were performed in accordance with the Declaration of Helsinki. Participants were all UNSW psychology undergraduate students and were recruited through the UNSW SONA system (Research management system) and received course credit.

##### Materials

Stimulus generation was performed in Matlab, with procedures illustrated in **Figure 2**. Each stimulus was a composite image, generated by morphing individual isoluminant masks of both younger and older faces onto the set of 160 individual greyscale images. Base and mask images were transformed from sRGB coordinated to the CIE L*a*b* space (CIE (International Commission on Illumination), 2004). Composite images were produced by combining the L* (luminance) component of the base image, and the a* and b* components of the relevant isoluminant mask. Isoluminant masks were based on individual images instead of an average image, to preserve the subtle color information of individual faces, such as small patches of uneven color distribution, which would be lost if an average image were used.

To spatially align an isoluminant mask with a target base image, we morphed the isoluminant mask to align 63 facial landmarks across the image pair (**Figure 2B**). These landmarks were adapted from the most common 68-point distribution model (Wang et al., 2018). To avoid potential visual artifacts between lips during image morphing, we used average landmarks between 5 pairs of corresponding points on the upper and lower lips, reducing the total number of landmarks from 68 to 63. In total, 320 images were generated, with each of the 160 base face images used with one younger and one older isoluminant mask of the same gender. Each isoluminant mask image was morphed onto 8 different base images, randomly selected. Lastly, we cropped the composite faces into an ellipse shape to exclude the hair and background, using the 63 landmarks to identify the facial area (**Figure 2C**).

##### Procedure

Stimuli were displayed on a 32” Display++ LCD Monitor by Cambridge Research Systems (resolution 1920×1080, refresh rate 120 Hz), using Windows (version 10) and Matlab (R2021a) with routines from Psychtoolbox 3 (Brainard, 1997; Kleiner et al., 2007; Pelli, 1997). Testing took place in darkened rooms, and participants viewed the displays from approximately 50 cm. The screen background was held at mean grey (luminance 60 cd/m^2^ throughout the experiment.

On each trial, a single, centrally-presented elliptical stimulus (7.4° in height, 5.7° in width) was displayed for 2 s then removed. A written prompt then asked the participant to rate the age of face by entering a two-digit integer (between 10 and 99) on the number pad of a computer keyboard. The prompt and text box would then remain on screen until the participant made their response. Participants were not informed about the true age range of the stimuli and no feedback was given on the accuracy of their responses. Each participant rated each of the 120 base faces only once, with 60 faces in the younger and 60 in the older isoluminant condition. Stimuli were counterbalanced such that, across participants, each base stimulus identity was presented the same number of times in the younger and older isoluminant conditions.

##### Data cleaning

Participants generally had high correlations between their age ratings and the true ages of the greyscale images (Pearson’s *R*, mean = 0.85, std = 0.09). We excluded data from 3 participants identified as outliers, whose correlations were greater than 3 scaled median absolute deviations from the median (*R* ≤ 0.65). Of the remaining data (8320 trials from 52 participants), we excluded data from individual trials with outlier values in the absolute difference between the true age and the rating (> 3 scaled median absolute deviations, or more than 24 years, total 326 trials excluded). Lastly, we excluded 12 trials where ratings were less than 14 years old, as all face images used were of adult faces. In total, 96% of the total trials were retained, and each stimulus had ratings from 16 to 27 participants (mean = 25.0, SD =1.5).

### Results

We tested whether facial age estimation is affected by color changes related to the aging process. Participants rated the age of a series of composite images, where a greyscale base image was paired with the isoluminant color from either a younger or older face. The distributions of rated ages are shown in **Figure 3**. We fit the average rated ages for each identity in both younger and older isoluminant conditions with a general linear model (with identity link function). A null model (intercept + true age, AIC = 1825.9, df = 318) was outperformed by a model including stimulus isoluminant condition (intercept + true age + isoluminant color; ΔAIC = -0.742, df = 317, X^2^=111.3, *p*=0.011), but this model fit was not further improved by adding an interaction term (intercept + true age + isoluminant color + true age * isoluminant color; ΔAIC = 1.34, df = 316, X^2^=11.2, *p*=0.419). The best-fitting model, depicted in **Figure 3A**, included a weighting term of perceived age being 1.18 years younger if the stimulus was shown in the younger, rather than the older, isoluminant condition. That is, on average, faces were judged as significantly younger when shown with the younger, rather than older, isoluminant color.

**Figure 3:**
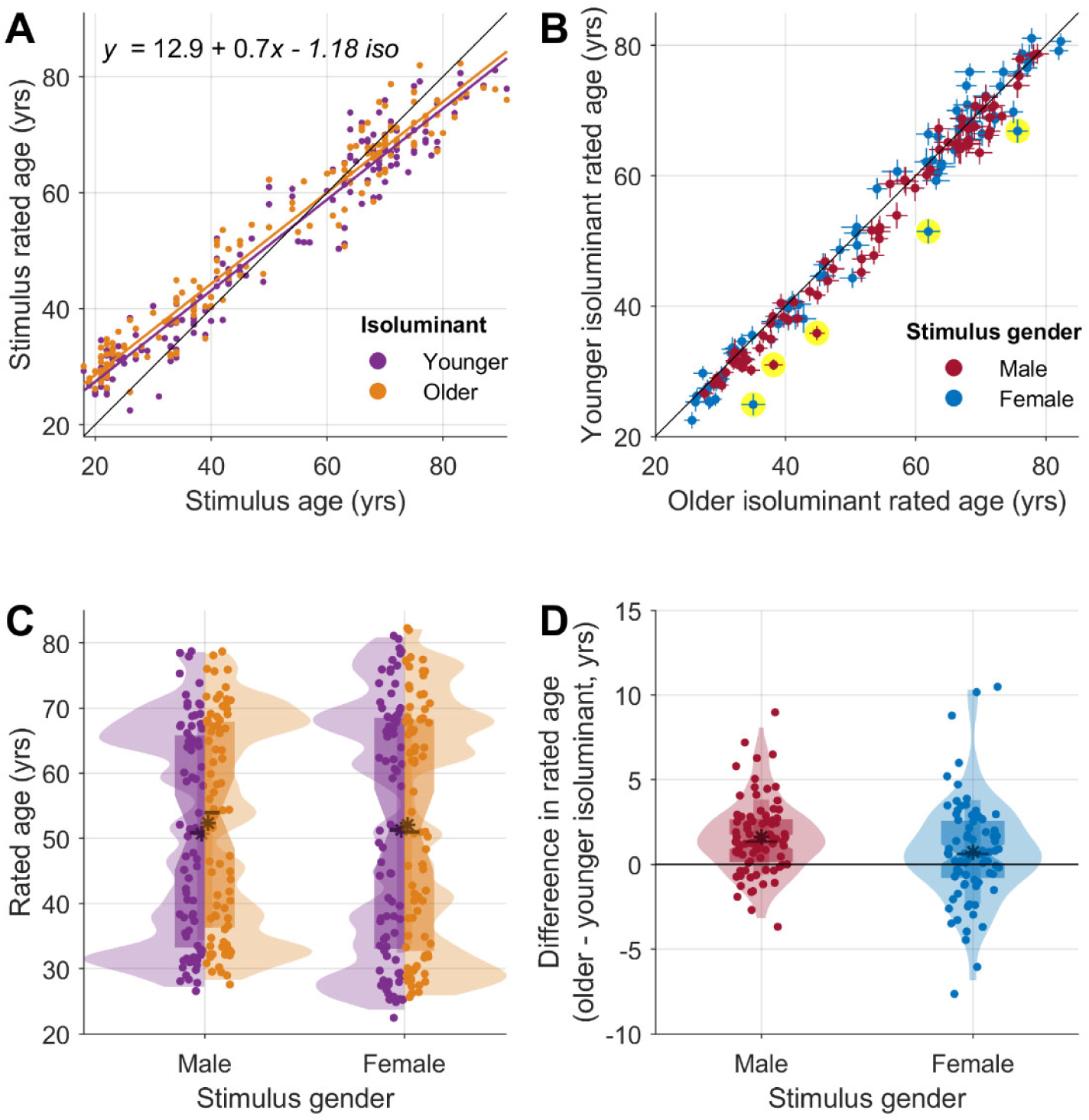
Age ratings (Experiment 1). Across all plots, each data point represents one of the 320 unique stimuli (**A** and **C**), or one of the 120 unique stimulus identities (**B** and **D**), and the black diagonal lines are reference lines of equality (**A** and **C**). **A**: The best-fitting model (see text for details) is shown with the orange and purple lines. **B**: Direct comparison of average rated ages for the two isoluminant conditions: points below the black line are images that have younger ratings in the young isoluminant condition compared to the old isoluminant condition. Error bars indicate +/-1 standard error of the mean. There were 5 cases where the with a significant difference in rated age across isoluminant conditions (q < 0.05), which are highlighted in yellow. **C**: Distributions of rated ages in younger (purple) and older (orange) isoluminant conditions, separated by stimulus gender. **D**: Rated age was significantly younger for both male and female stimuli, but this difference was significantly greater for male faces (see text for statistical tests).

To test how this shift varied across individual image identities, we directly compared age ratings in the younger and older isoluminant conditions for each identity (**Figure 3B**). In most cases, average ratings were younger when the identity was shown with the younger isoluminant color (i.e. most points are below the diagonal in **Figure 3B**). Paired *t*-tests, with FDR correction for multiple comparisons (Benjamini & Hochberg, 1995), revealed 5 identities where average ratings were significantly different (q<0.05), in each case in the direction of the younger isoluminant condition being rated as younger.

Next, we considered whether the effect of isoluminant color on perceived age varied across male and female image identities (**Figure 3C-D**). When data from male and female faces were considered individually with Wilcoxon tests, the median perceived age difference was significantly different from 0 for male faces (median difference = 1.34 years; *W* = 2736, z= 5.65 *p*<0.001), and for female faces (median difference = 0.62 years; *W* = 2095, z = 2.28, *p* = 0.023). A Mann-Whitney U-test for equal medians showed that the effect of gender on perceived age difference was of significantly larger magnitude for male faces than for female faces, *U* = 5796, z = 2.196, *p* = 0.028.

#### Experiment 2

##### Overview

Previously, we found that faces were perceived as more youthful when RG contrast (and overall redness) is increased (Hsieh et al., 2023); however, in the sample of faces used here, average redness (a* value) increases with age in the original images (**Figure 1**), which predicts an effect between redness and perceived age in the opposite direction. A limitation of our previous work (Hsieh et al., 2023) was that the model ages were not included with the stimulus images, and all selected images used were limited to between 20 and 40 years old based on the subjective age ratings provided, meaning that the findings may not generalize to other age groups (Voegeli et al., 2021).

In Experiment 2 we used the faces used in Experiment 1, which had self-reported ages of a wide range (18-91 years), and measured the effect of varying RG contrast on perceived youthfulness. Participants viewed each face with varying redness levels, generated by scaling the contrast of the CIE a* component. They were asked to select the one that appeared youngest for a passport photo, with an implicit consideration of the apparent ’naturalness’ of the redness variation. Research has shown that people prefer facial color within the normal range, with very high levels of redness being associated with increased perceived aggression but decreased attractiveness (Stephen et al., 2012). We aimed to determine whether the optimal redness level that makes faces appear the youngest would be significantly redder than the originals and whether the optimal redness level for youthfulness would vary across different age groups. Based on our previous work (Hsieh et al., 2023), we hypothesized that the optimal redness level to make faces look the youngest would be significantly redder than the originals.

#### Methods

##### Participants

A separate sample of 55 participants completed Experiment 2, with 22 male and 33 female participants, aged 17-53 years (mean = 19.7, SD = 5.72). All participants had normal or corrected to normal vision and normal color vision (self-reported). All consent and recruitment procedures were as in Experiment 1.

##### Materials

Stimuli for Experiment 2 were based on the same sample of 160 images that were used in Experiment 1 (80 male, 80 female, ranging in age from 18 to 91). We used multiplicative scaling of the a* content (the red-green dimension of CIE L*a*b* space) of each original image to experimentally adjust redness level of the images. We used 9 different multiplicative scaling factors (*m*), defined as *m* = 2*^x^*, with 9 values of *x*, linearly spaced between -0.8 and 0.8, giving values of *m* ranging from 0.57 to 1.74 with logarithmic spacing. This range was slightly greater than the range of multiplication factors used in a previous study with the same stimulus manipulation (Hsieh et al., 2023). We used *x* to denote RG contrast factors in our data analysis, since values of *x* were centred on 0 and had equal spacing, ensuring an average response of 0 if participants were to give random answers. Example stimuli are shown in **Figure 4**.

**Figure 4:**
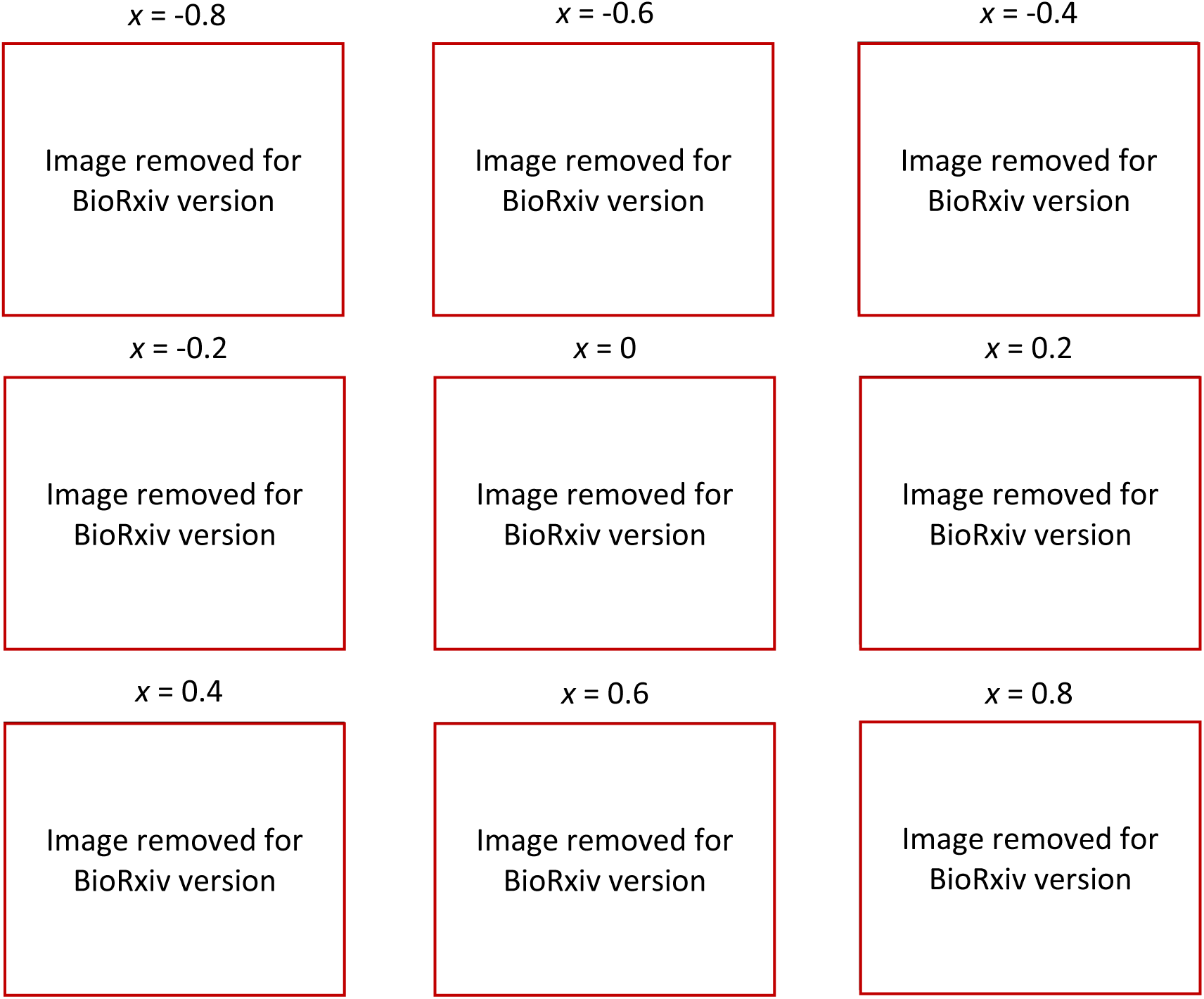
Experiment 2 stimuli. Images were manipulated by RG contrast factors (*x*) indicated above. In the experiment, the positions of images were randomized, and the numbers were not shown.

##### Procedure

Each participant completed 160 trials. On each trial, a set of 9 images of the same face with varying RG contrast factors (**Figure 4**) positioned randomly in a 3×3 array was presented for 1 s, with positions numbered 1-9. The images then disappeared, and a written prompt instructed participants to: *’Please help your friend pick a passport photo that makes them look as young as possible*.’ We used this naturalistic task to encourage an implicit consideration of whether the redness was within the typical range of redness variation, without complicating the task by adding an explicit ‘naturalness’ constraint. Participants responded using the corresponding key on the number pad (1 to 9) followed by ‘enter’ to submit their response. The prompt and response options remained visible until they responded. A total of 8,800 ratings were collected from the 55 participants.

##### Data cleaning

No participants or trials were excluded as there is no reliable method to exclude responses given the purely subjective nature of the judgement.

### Results

We tested which RG contrast factor was judged as optimal for maximising perceived youthfulness. Results are shown in **Figure 5**. On average, participants selected images with positive RG contrast factors (**Figure 5A**), with a mean RG contrast factor (*x*) of 0.17, which corresponds to a 12.5% increase in the a* contrast relative to the original image. When selected RG contrast factors for each image identity were averaged, the averages were above zero in almost all cases (**Figure 5B**). One-sample *t*-tests revealed that these averages were significantly different from the original, unfiltered image (RG contrast factor = 0), for both male (*t*(159) = 11.81, *d* = 1.32, *p* < 0.001) and female faces (*t*(159) = 12.13, *d* = 1.36 *p* < 0.001). The average RG contrast was slightly higher for female faces (mean = 0.19, or 14.1% increase) than for male faces (mean = 0.15, or 11.0% increase): a two-sample t-test showed that this difference approached significance (*t*(158) = 1.94, *d* = 0.307, *p* = 0.054).

**Figure 5:**
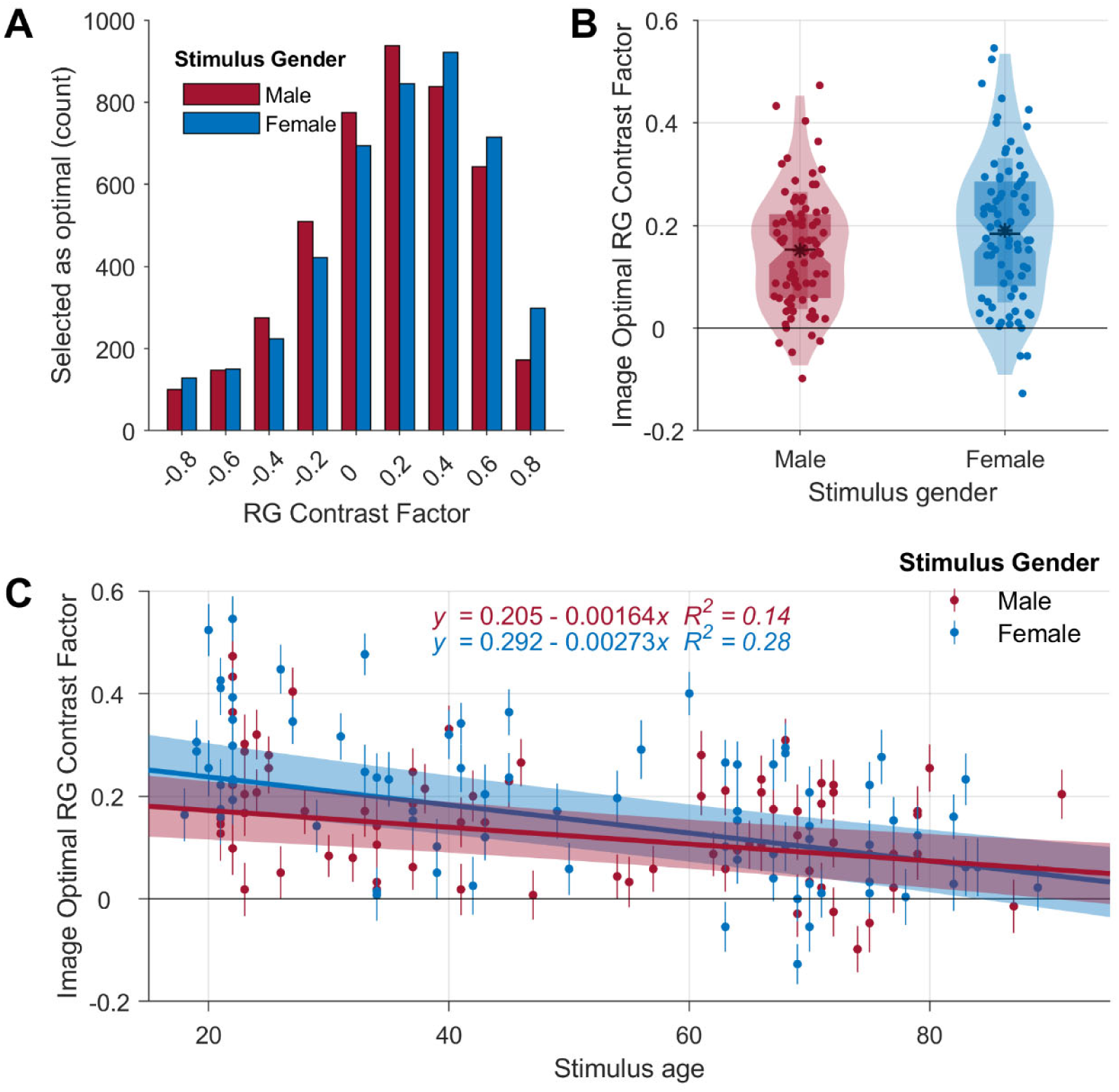
Optimal RG contrast (Experiment 2). **A**: Distributions of selected RG contrast factors across all 8,800 trials. **B**: Mean RG contrast factor by stimulus image. Each point (n=160) indicates the RG contrast factor selected for each stimulus image, averaged across ratings from 55 participants. **C**: Mean RG contrast response as a function of true stimulus age. Each of the 160 data points represents a particular stimulus identity. Formulae are for the fitted trend lines (for male and female faces separately), with shaded areas showing 95% confidence intervals.

We tested how the selected RG factor varied with stimulus true age, we found that true age significantly influenced the RG contrast factor, *R^2^*= .20, *t*(158) = -6.35, β = -.002, *p* < .001, 95% CI [-0.0028, -0.0015]. β is the slope of the regression line and is significantly different from zero. **Figure 5C** shows these regression lines separately for male and female faces; in both cases, the optimal RG contrast factor decreases with age, and females tend to require higher redness level compared to males to make them look the youngest.

## Discussion

These experiments aimed to answer the following questions: do age-related color changes affect perceived age independently of luminance information? Would the optimal redness level to make faces appear the youngest be significantly redder than the originals across a broad range of ages? Our findings demonstrate that age-related color changes do provide cues for age perception, dissociable from luminance information. We also found that the optimal redness level for maximising apparent youthfulness was significantly redder than the originals across a broad range of ages, but that for images of younger individuals, the optimal level was at a greater level of accentuated redness than for their older counterparts. Additionally, we found that females require a higher redness compared to males to make them appear the youngest.

Experiment 1 establishes that age-related color changes provide cues that affect perceived age. The color changes employed were purely chromatic, distinguishing them from previous studies that included both chromatic and luminance changes (Burt & Perrett, 1995; Fink et al., 2006). Our analysis of simple statistics of a* and b* distributions within each image showed that age tends to be positively correlated with average a* and b* values. If participants used the positive correlation with average redness as a cue to age, then increasing redness should increase perceived age, as found by Lu et al. (2021) for unmanipulated images of younger adults, but in contrast to the effect found by Voegeli et al. (2023). Furthermore, average redness as a cue cannot account for the effect in the opposite direction we found previously (Hsieh et al., 2023), which was replicated here in Experiment 2. This suggests that the effect of color on perceived age in Experiment 1 is driven by spatially specific changes in color distribution, such as chromatic cues to skin evenness, which is negatively correlated with chronological and perceived age (Fink, Bunse, et al., 2012; Gunn et al., 2009; Leyden, 1990; Matts et al., 2007; Nkengne et al., 2008).

Color changes within specific facial regions or features may also drive changes in perceived age. Some facial features may carry greater weight in age estimation tasks, such as the under-eye region, or the presence of skin wrinkles (Gunn et al., 2009; Messaraa et al., 2020), but these effects may be driven by luminance information, meaning that they are not necessarily the features where chromatic information has the greatest effect on perceived age. The lips are a facial feature where chromatic information has a demonstrated effect on perceived age: Porcheron et al., (2013) report that local a* value within the lips decreases with age, and that increasing the redness of the lips significantly reduces perceived age. Messaraa et al. (2020) also report that lip redness (a*) is negatively correlated with perceived age, but only within older adults (>50 years), with no correlation for younger adults (<50 years). Overall, our results demonstrate that chromatic information contributes to perceived age independently of luminance information, but further work is needed to determine the how specific spatial-chromatic properties affect perceived age.

In Experiment 2, when participants were asked to select face images perceived as youngest, they selected images where RG contrast (a* contrast) was increased by approximately 10.2% compared with the original. Previously, we found that increasing RG contrast by 50% research led to images being rated as significantly younger than those same images where RG contrast was decreased by 50% (Hsieh et al., 2023), for stimuli limited to adults 20-40 years old based on the subjective age ratings. The average RG contrasts obtained in Experiment 2 suggest that the manipulations in previous research may have been excessive for many images, although there was substantial variability across images, and there were individual images where the average optimal RG contrast approached a 50% increase (**Figure 5**). While participants were instructed to choose the “youngest” looking image, our design did not measure how many years younger the selected redness levels made the original image appear, though in our previous work the average difference in rated age between RG decreased and increased faces was less than a year (approximately 7 months) (Hsieh et al., 2023).

The optimal RG contrast for appearing youngest varied both with gender and with age. The optimal contrast was higher overall for female than for male faces, and both the optimal contrast and the gender difference decreased as age increased. Based on the lines of best fit (**Figure 5**), for 20 year olds, the optimal RG contrasts were 17.9% (female faces) and 12.7% (male faces). By 80 years old, the optimal RG contrast remained above zero, but were lower (5.2% for female faces; 5.3% for male faces). In a previous study, Stephen et al. (2009) asked participants to adjust the RG contrast of facial skin, excluding lips, to maximize apparent health. Similar to our results for perceived age, Stephen et al. (2009) found that increasing RG contrast significantly enhanced perceived health, consistent with perceived age and perceived health being dissociable but correlated measures (Fink et al., 2018; Fink, Matts, et al., 2012; Messaraa et al., 2020). However, in contrast to our gender effect, they found that male faces required a significantly higher redness level than female faces to appear the healthiest (Stephen et al., 2009). This may reflect a dissociation between perceptions of age and health, or the gender effects observed may be specific to the set of face stimuli used. Future studies where participants provide age and health ratings for the same set of RG contrast manipulated face stimuli could resolve whether these gender-based differences are common to perceived age and perceived health or if this reflects a dissociation between perceptions of these attributes.

Relatedly, the optimal redness levels we identified in Experiment 2 may be specific to age perception tasks. Despite the positive association between apparent youth and health, observers may prioritize different facial cues when estimating these characteristics (Fink et al., 2018; Fink & Matts, 2008; Messaraa et al., 2020). In age estimation tasks, skin surface topography appears more crucial than skin color distribution, while in health perception, skin color serves as a stronger cue (Fink et al., 2018; Fink & Matts, 2008). Given the differing weights of color in age and health perception, the optimal redness levels for making faces appear the youngest and healthiest may not be the same. Further investigations of the optimal redness levels for different facial attributes could give new insight into how chromatic information contributes to face perception.

Consistent with the results of Experiment 1, the change in optimal contrast with age suggests that isoluminant red-green information contributes to perceived age in a way beyond that captured by simple statistics such as average redness. For instance, if there are patterns of redness across the face that change reliably with age, then increasing the contrast of these signals for younger faces could accentuate their perceived youth, whereas for older faces increasing the visibility of patterns of redness might accentuate not only signals associated with youth (e.g. redness of the lips), but also signals associated with older age (e.g. skin heterogeneity). Overall, our results provide a clear demonstration that chromatic information makes a significant contribution to the perceived age of faces, independent of luminance information, and suggests that the spatial distribution of redness contributes to apparent youthfulness, especially in younger faces.

## Acknowledgements

This work was supported by Australian Research Council Discovery Project DP220100747 to EG.

## References

1. Benjamini, Y., & Hochberg, Y. (1995). Controlling the False Discovery Rate: A Practical and Powerful Approach to Multiple Testing. Journal of the Royal Statistical Society: Series B (Methodological), 57(1), 289–300. 10.1111/j.2517-6161.1995.tb02031.x

2. Brainard, D. H. (1997). The Psychophysics Toolbox. Spatial Vision, 10(4), 433–436. 10.1163/156856897x00357

3. Bruce, V., & Langton, S. (1994). The Use of Pigmentation and Shading Information in Recognising the Sex and Identities of Faces. Perception, 23(7), 803–822. 10.1068/p230803

4. Burt, D. M., & Perrett, D. I. (1995). Perception of age in adult Caucasian male faces: Computer graphic manipulation of shape and color information. Proceedings of the Royal Society of London. Series B: Biological Sciences, 259(1355), 137–143. 10.1098/rspb.1995.0021

5. CIE (International Commission on Illumination). (2004). Colorimetry, 3rd edition (CIE 015:2004). https://cie.co.at/publications/colorimetry-3rd-edition

6. Dobos, G., Trojahn, C., Lichterfeld, A., D′Alessandro, B., Patwardhan, S. V., Canfield, D., Blume-Peytavi, U., & Kottner, J. (2015). Quantifying dyspigmentation in facial skin ageing: An explorative study. International Journal of Cosmetic Science, 37(5), 542–549. 10.1111/ics.12233

7. Fink, B., Bunse, L., Matts, P. J., & D’Emiliano, D. (2012). Visible skin coloration predicts perception of male facial age, health and attractiveness. International Journal of Cosmetic Science, 34(4), 307–310. 10.1111/j.1468-2494.2012.00724.x

8. Fink, B., Grammer, K., & Matts, P. J. (2006). Visible skin color distribution plays a role in the perception of age, attractiveness, and health in female faces. Evolution and Human Behavior, 27(6), 433–442. 10.1016/j.evolhumbehav.2006.08.007

9. Fink, B., & Matts, P. (2008). The effects of skin color distribution and topography cues on the perception of female facial age and health. Journal of the European Academy of Dermatology and Venereology, 22(4), 493–498. 10.1111/j.1468-3083.2007.02512.x

10. Fink, B., Matts, P. J., Brauckmann, C., & Gundlach, S. (2018). The effect of skin surface topography and skin coloration cues on perception of male facial age, health and attractiveness. International Journal of Cosmetic Science, 40(2), 193–198. 10.1111/ics.12451

11. Fink, B., Matts, P. j., D’Emiliano, D., Bunse, L., Weege, B., & Röder, S. (2012). Color homogeneity and visual perception of age, health and attractiveness of male facial skin. Journal of the European Academy of Dermatology and Venereology, 26(12), 1486–1492. 10.1111/j.1468-3083.2011.04316.x

12. George, P. A., & Hole, G. J. (2000). The role of spatial and surface cues in the age-processing of unfamiliar faces. Visual Cognition, 7(4), 485–509. 10.1080/135062800394621

13. Gomi, T., & Imamura, T. (2019). Age-related changes in the vasculature of the dermis of the upper lip vermilion. Aging (Albany NY), 11(11), 3551–3560. 10.18632/aging.101996

14. Gunn, D. A., Rexbye, H., Griffiths, C. E. M., Murray, P. G., Fereday, A., Catt, S. D., Tomlin, C. C., Strongitharm, B. H., Perrett, D. I., Catt, M., Mayes, A. E., Messenger, A. G., Green, M. R., Ouderaa, F. van der, Vaupel, J. W., & Christensen, K. (2009). Why Some Women Look Young for Their Age. PLOS ONE, 4(12), e8021. 10.1371/journal.pone.0008021

15. Hill, H., Bruce, V., & Akamatsu, S. (1995). Perceiving the sex and race of faces: The role of shape and color. Proceedings of the Royal Society of London. Series B: Biological Sciences, 261(1362), 367–373. 10.1098/rspb.1995.0161

16. Hsieh, J. Y. J., Boyce, W. P., Goddard, E., & Clifford, C. W. G. (2023). Color information biases facial age estimation and reduces inter-observer variability. Scientific Reports, 13(1), 13224. 10.1038/s41598-023-39902-z

17. Kemp, R., Pike, G., White, P., & Musselman, A. (1996). Perception and Recognition of Normal and Negative Faces: The Role of Shape from Shading and Pigmentation Cues. Perception, 25(1), 37–52. 10.1068/p250037

18. Kikuchi, K., Katsuyama, M., Yoshikawa, T., Wakeshima, N., & Iizuka, H. (2022). Epidemiological survey of the quantity and anatomical position of facial pigmented spots in relation to age in 454 Japanese women. The Journal of Dermatology, 49(12), 1245–1254. 10.1111/1346-8138.16561

19. Kleiner, M., Brainard, D., & Pelli, D. G. (2007). What’s new in Psychtoolbox-3? Perception, 36(ECVP Abstract Supplement), 14. 10.1177/03010066070360S101

20. Leyden, J. J. (1990). Clinical features of ageing skin. British Journal of Dermatology, 122(s35), 1–3. 10.1111/j.1365-2133.1990.tb16118.x

21. Lu, Y., Yang, J., Xiao, K., Pointer, M., Li, C., & Wuerger, S. (2021). Skin coloration is a culturally-specific cue for attractiveness, healthiness, and youthfulness in observers of Chinese and western European descent. PLOS ONE, 16(10), e0259276. 10.1371/journal.pone.0259276

22. Matts, P. J., Fink, B., Grammer, K., & Burquest, M. (2007). Color homogeneity and visual perception of age, health, and attractiveness of female facial skin. Journal of the American Academy of Dermatology, 57(6), 977–984. 10.1016/j.jaad.2007.07.040

23. Mayes, A., Murray, P. G., Gunn, D. A., Tomlin, C. C., Catt, S. D., Wen, Y. B., Zhou, L. P., Wang, H. Q., Catt, M., & Granger, S. P. (2010a). Environmental and Lifestyle Factors Associated with Perceived Facial Age in Chinese Women. PLOS ONE, 5(12), e15270. 10.1371/journal.pone.0015270

24. Mayes, A., Murray, P., Gunn, D., Tomlin, C., Catt, S., Wen, Y., Zhou, L., Wang, H., Catt, M., & Granger, S. (2010b). Ageing appearance in China: Biophysical profile of facial skin and its relationship to perceived age. Journal of the European Academy of Dermatology and Venereology, 24(3), 341–348. 10.1111/j.1468-3083.2009.03418.x

25. Messaraa, C., Richard, T. J. C., Walsh, M., Doyle, L., O’Connor, C., Robertson, N., Mansfield, A., Hurley, S., Mavon, A., & Grenz, A. (2020). Perceived age and perceived health among a Chinese cohort: Does it mean the same thing? International Journal of Cosmetic Science, 42(5), 471–481. 10.1111/ics.12647

26. Minear, M., & Park, D. C. (2004). A lifespan database of adult facial stimuli. Behavior Research Methods, Instruments, & Computers, 36(4), 630–633. 10.3758/BF03206543

27. Nkengne, A., Bertin, C., Stamatas, G., Giron, A., Rossi, A., Issachar, N., & Fertil, B. (2008). Influence of facial skin attributes on the perceived age of Caucasian women. Journal of the European Academy of Dermatology and Venereology, 22(8), 982–991. 10.1111/j.1468-3083.2008.02698.x

28. Ortonne, J.-P. (1990). Pigmentary changes of the ageing skin. British Journal of Dermatology, 122(s35), 21–28. 10.1111/j.1365-2133.1990.tb16121.x

29. Palermo, R., & Rhodes, G. (2007). Are you always on my mind? A review of how face perception and attention interact. *Neuropsychologia*, The Perception of Emotion and Social Cues in Faces, 45(1), 75–92. 10.1016/j.neuropsychologia.2006.04.025

30. Pelli, D. G. (1997). The VideoToolbox software for visual psychophysics: Transforming numbers into movies. Spatial Vision, 10(4), 437–442. 10.1163/156856897x00366

31. Porcheron, A., Mauger, E., & Russell, R. (2013). Aspects of Facial Contrast Decrease with Age and Are Cues for Age Perception. PLOS ONE, 8(3), e57985. 10.1371/journal.pone.0057985

32. Rhodes, G. (2009). Age estimation of faces: A review. Applied Cognitive Psychology, 23(1), 1–12. 10.1002/acp.1442

33. Russell, R. (2003). Sex, Beauty, and the Relative Luminance of Facial Features. Perception, 32(9), 1093–1107. 10.1068/p5101

34. Russell, R., Kramer, S. S., & Jones, A. L. (2017). Facial Contrast Declines with Age but Remains Sexually Dimorphic Throughout Adulthood. Adaptive Human Behavior and Physiology, 3(4), 293–303. 10.1007/s40750-017-0068-x

35. Russell, R., Sinha, P., Biederman, I., & Nederhouser, M. (2006). Is Pigmentation Important for Face Recognition? Evidence from Contrast Negation. Perception, 35(6), 749–759. 10.1068/p5490

36. Russell, R., Sweda, J. R., Porcheron, A., & Mauger, E. (2014). Sclera color changes with age and is a cue for perceiving age, health, and beauty. Psychology and Aging, 29(3), 626–635. 10.1037/a0036142

37. Sforza, C., Grandi, G., De Menezes, M., Tartaglia, G. M., & Ferrario, V. F. (2011). Age- and sex-related changes in the normal human external nose. Forensic Science International, 204(1), 205.e1-205.e9. 10.1016/j.forsciint.2010.07.027

38. Stephen, I. D., Law Smith, M. J., Stirrat, M. R., & Perrett, D. I. (2009). Facial Skin Coloration Affects Perceived Health of Human Faces. International Journal of Primatology, 30(6), 845–857. 10.1007/s10764-009-9380-z

39. Stephen, I. D., Oldham, F. H., Perrett, D. I., & Barton, R. A. (2012). Redness enhances perceived aggression, dominance and attractiveness in men’s faces. Evolutionary Psychology, 10(3), 562–572. Scopus. 10.1177/147470491201000312

40. Voegeli, R., Campiche, R., Biassin, R., Rawlings, A. V., Shackelford, T. K., & Fink, B. (2023). Predictors of female age, health and attractiveness perception from skin feature analysis of digital portraits in five ethnic groups. International Journal of Cosmetic Science, 45(5), 672–687. 10.1111/ics.12877

41. Voegeli, R., Schoop, R., Prestat-Marquis, E., Rawlings, A. V., Shackelford, T. K., & Fink, B. (2021). Differences between perceived age and chronological age in women: A multi-ethnic and multi-centre study. International Journal of Cosmetic Science, 43(5), 547–560. 10.1111/ics.12727

42. Wang, N., Gao, X., Tao, D., Yang, H., & Li, X. (2018). Facial feature point detection: A comprehensive survey. Neurocomputing, 275, 50–65. 10.1016/j.neucom.2017.05.013

43. Willner, P., & Rowe, G. (2001). Alcohol servers’ estimates of young people’s ages. *Drugs: Education*, Prevention & Policy, 8(4), 375–383. 10.1080/09687630010019299

